# Variation in temperature but not diet determines the stability of latitudinal clines in tolerance traits and their plasticity

**DOI:** 10.1101/2025.05.20.655196

**Authors:** Greg M. Walter, Avishikta Chakraborty, Fiona E. Cockerell, Vanessa Kellermann, Matthew D. Hall, Craig R. White, Carla M. Sgrò

## Abstract

Latitudinal clines are routinely used as evidence of adaptation across broad climatic gradients. However, if environmental variation influences the strength of latitudinal clines, then clinal patterns will be unstable, and using patterns of adaptation to predict population responses to global change will be difficult. To test whether environmental variation influences latitudinal clines, we sampled five populations of *Drosophila melanogaster* spanning 3000-km of east coast Australia, and measured stress tolerance (heat, cold and desiccation) and body size on flies that developed in six combinations of temperature (13°C, 25°C and 29°C) and diet (standard and low-calorie) treatments. We found latitudinal clines where populations further from the equator had larger wings, higher cold tolerance and lower heat tolerance. For all traits, temperature determined the strength of latitudinal clines, whereas diet had little influence. Steeper clines often emerged in warmer treatments, created by latitudinal clines in plasticity. In the warmest temperature, higher latitude populations showed larger increases in heat tolerance, larger reductions in desiccation tolerance but smaller decreases in cold tolerance. Heat tolerance was the only trait that supported the climate variability hypothesis and a trade-off between plasticity and tolerance. Environment-dependent latitudinal clines are therefore likely to determine variation in population responses to global change.

## Introduction

Determining how natural populations adapt to environmental variation across geography is crucial if we are to accurately predict their response to ongoing global change (Hoffmann and Sgrò 2011; Sgrò et al. 2011; Valladares et al. 2014). Latitudinal clines in ecologically important traits emerge in response to local adaptation across large climatic gradients (Huey et al. 2000; Hoffmann et al. 2002; Etterson 2004; Trotta et al. 2006; Arthur et al. 2008; Hopkins et al. 2008; Hernández et al. 2019; Lush et al. 2024) and are widely used to predict population responses to future environmental change (Hoffmann and Sgrò 2011; Sunday et al. 2011; Mayekar et al. 2022). However, current predictions of population responses to global change often assume that latitudinal patterns of adaptation are stable. For example, populations from warmer environments that have evolved higher heat tolerance are generally expected to respond better to warmer conditions created by climate change (Lush et al. 2024). If, however, populations along latitude vary in their response to environmental variation, then predicting population responses to global change will need to consider how populations vary in their sensitivity to the environment (Chevin et al. 2013; Valladares et al. 2014; Matesanz and Ramírez-Valiente 2019; Walter et al. 2020). While studies have emphasised the importance of incorporating local adaptation in predictive models to increase the accuracy of predictions under climate change (Chevin et al. 2010; Reed et al. 2011; Bennett et al. 2019; Meek et al. 2023), we still lack a fundamental understanding of how patterns of adaptation change in response to environmental variation.

Latitudinal clines are typically estimated under a single temperature in common garden conditions, meaning that they rarely consider whether latitudinal patterns remain stable as the environment changes (Liefting et al. 2009). Testing whether patterns of adaptation, such as along a latitudinal gradient, remain stable across environments can reveal the extent to which locally adapted populations are likely to vary in their response to global change (DeMarche et al. 2018; DeMarche et al. 2019). While latitudinal clines in climatic stress tolerance traits can remain stable across sexes (Lasne et al. 2018) and simulated seasonal changes (Lush et al. 2024), pathogen infection can erase latitudinal clines in thermal tolerance (Hector et al. 2020), and patterns of climatic adaptation across latitudinal extremes can change with temperature (Liefting et al. 2009; van Heerwaarden and Sgrò 2011) and combinations of temperature and diet (Alton et al. 2020; Chakraborty et al. 2020). This suggests that variation in the environment could determine the strength of latitudinal clines that represent adaptation across a broad climatic gradient. However, we lack an understanding of how combinations of environmental stressors determine the stability of latitudinal clines in climate tolerance, which limits our ability to accurately predict population responses to climate change (Louthan et al. 2021; Zettlemoyer 2023).

The capacity for individuals to adjust their phenotype to cope with environmental variation, known as phenotypic plasticity, is expected to help populations cope with environmental variation created by climate change (Via et al. 1995; Ghalambor et al. 2007; Charmantier et al. 2008; Reed et al. 2011; Morris 2014; Anderson and Gezon 2015; Acasuso-Rivero et al. 2019). While we know that adaptation to different environments can create changes in plasticity (Murren et al. 2014; Matesanz and Ramírez-Valiente 2019; Chakraborty et al. 2020; Walter et al. 2022), we still have a relatively poor understanding of how environmental heterogeneity generates plasticity. Two main theories are expected to underlie the evolution of plasticity: the climate variability hypothesis, and the plasticity trade-off hypothesis. Under the climate variability hypothesis, greater temperature heterogeneity is expected to increase thermal plasticity (Janzen 1967; Ghalambor et al. 2006). Alternatively, greater plasticity might also trade-off with overall thermal tolerance because high thermal tolerance should not be possible alongside high plasticity (Roff and Fairbairn 2007; van Heerwaarden and Kellermann 2020). Across latitude, these two processes should reinforce each other for upper thermal limits because constant high temperatures at lower latitudes will favour higher heat tolerance and lower plasticity. However, it remains unclear whether these same patterns will emerge for other traits, such as lower thermal limits, because variable cooler temperatures at higher latitudes should select for both colder tolerance and higher plasticity. Understanding how locally adapted populations along latitude evolve differences in thermal tolerance and plasticity for multiple traits can reveal how populations adapt to environmental heterogeneity, which will help predict their response to climate change.

Supporting the climate variability hypothesis, ectotherm species from higher latitudes show greater capacity to thermally acclimate (Seebacher et al. 2015; Kellermann and van Heerwaarden 2019) and have broader thermal tolerances (Sunday et al. 2011). However, these studies focused on differences among species, which are less informative for predicting population responses (within species) to climate change (Meek et al. 2023). Studies comparing populations of the same species found no latitudinal patterns in plasticity (van Heerwaarden et al. 2014; Chakraborty et al. 2020), or the opposite pattern where low latitude populations showed greater plastic acclimation after a hardening treatment (Cockerell et al. 2014). The strongest evidence that plasticity increases in populations from high latitudes is for a variety of fitness and foliar traits in plants (Molina-Montenegro and Naya 2012; Ren et al. 2020), whereas studies on ectotherms only focus on populations from latitudinal extremes (Mathur and Schmidt 2017; van Heerwaarden and Sgrò 2017; Cicchino et al. 2024). In contrast to the wealth of studies that test the climate variability hypothesis, surprisingly few have tested for a trade-off between plasticity and overall thermal tolerance, which means that evidence of a trade-off is scarce (Gunderson and Stillman 2015; van Heerwaarden et al. 2016; Gunderson 2023). While a trade-off for desiccation resistance was found in *Drosophila*, there was no relationship with latitude (Kellermann and van Heerwaarden 2025). To more accurately predict how populations will respond to global change, further manipulative experiments are therefore required to test whether adaptation across latitude is associated with clinal patterns in thermal plasticity and the presence of a trade-off with overall thermal tolerance (Kellermann and van Heerwaarden 2019).

To test whether latitudinal clines in tolerance traits remain stable under combinations of diet and temperature treatments, we studied populations from a well-established latitudinal cline of *Drosophila melanogaster* along east-coast Australia (Hoffmann et al. 2002; Hoffmann and Weeks 2007; Sgrò et al. 2010). We used a manipulative laboratory experiment to test how clinal patterns in body size and tolerance traits, and their plasticity, changed across combinations of diet and temperature treatments. We sampled five populations of *D. melanogaster* across a 3000-km latitudinal cline in eastern Australia from Tasmania (43°S) to North Queensland (18°S; **Fig. 1**). While temperature decreases at higher latitudes (**Fig. 1**), variation in temperature tends to increase and rainfall decreases (**Table S1**). Latitudinal clines therefore represent adaptation to a combination of climate variables. In the lab, we reared larvae from each population in six treatments (three temperatures × two diets), and measured tolerance traits (heat, cold and desiccation) and wing size on the adults that emerged. We tested whether combinations of temperature and diet treatments determined the strength and directionality of latitudinal clines in trait means and their plasticity (estimated across treatments). Supporting the climate variability hypothesis and the trade-off between basal tolerance and plasticity, we predicted that populations from higher latitudes would show greater plasticity but lower basal tolerance in thermal tolerance traits when compared to low latitude populations.

**Fig. 1.**
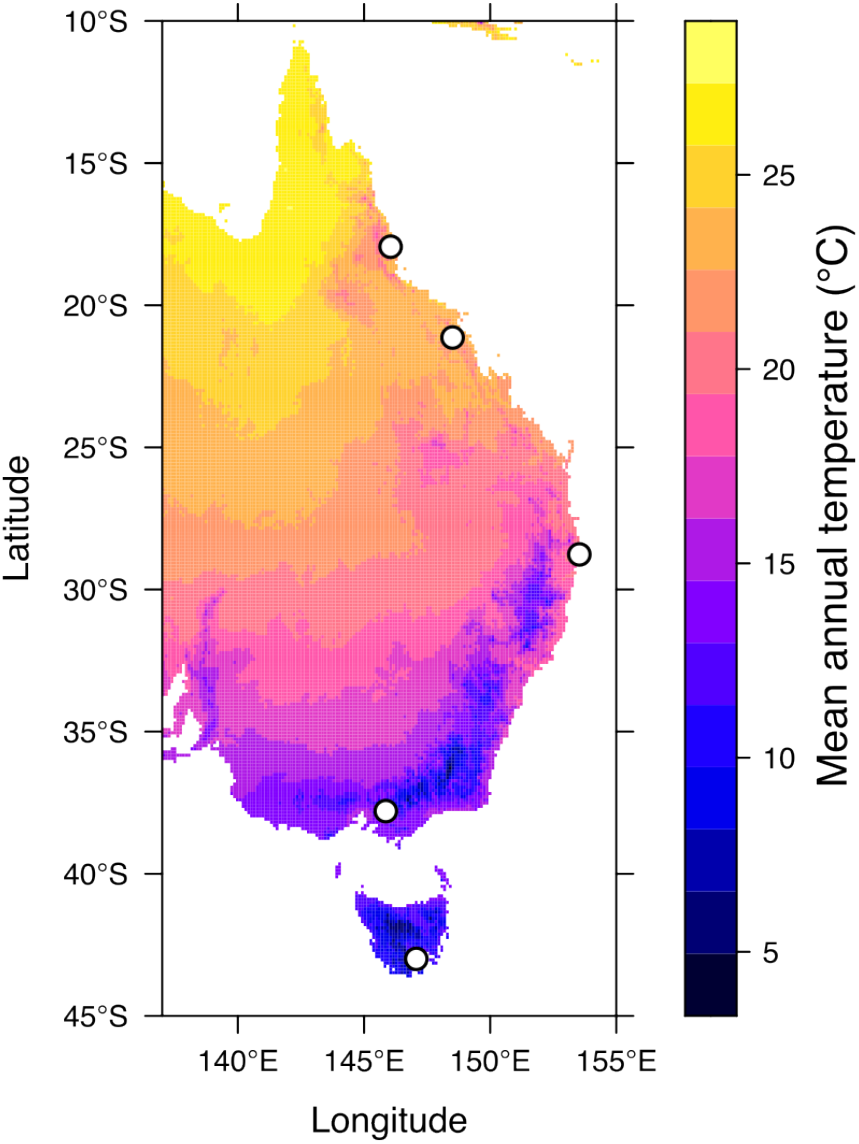
Map of sampling locations along the cline (n=5; unfilled circles). Colour represents the average mean annual temperature. **Table S1** contains temperature variability and mean annual rainfall for each location.

## Methods

### Collection and maintenance of experimental populations

Between March and May 2021, we collected 20-50 field-inseminated female *Drosophila melanogaster* using sweep nets over fruit waste from fruit farms (primarily banana and apple) at five locations across Tasmania, New South Wales, Victoria, and Northern Queensland (**Fig. 1**; **Table S1**). In the laboratory, we established isofemale lines by allowing the collected females to lay eggs for four days in 40mL vials containing 7mL of a standard yeast-potato-dextrose medium (potato flakes 18.2g/L; dextrose 27.3g/L; Brewer’s yeast 36.4g/L; agar 6.4g/L, Nipagin [10% w/v in ethanol] 10.9mL/L; and propionic acid 4.6mL/L). The larvae were reared at 25°C on a 12:12 h light:dark cycle. In the subsequent generation, we verified species identification to avoid contamination from the morphologically similar *D. simulans*. In the third generation, we treated each isofemale line with 0.3mg/mL of tetracycline in the food to eliminate differences in endosymbionts, particularly *Wolbachia*, which is more common in tropical populations (Hoffmann 1988). After three generations, we combined five males with five virgin females from each isofemale line (19-47 lines per population, mean=35.8; **Table S1**) into a mass-bred population maintained in five 300mL bottles containing 62.5mL of standard food. Each generation, adults were collected and kept in bottles until they reached peak fertility (approximately four days), after which they were allowed to lay 250-300 eggs in each of the five bottles. After laying, adults were removed, pupation cards were added, and larvae were left to develop. We maintained the populations at 1000-1500 individuals at 25°C (12:12h light cycle) for 12 generations (approximately nine months) to ensure sufficient mixing of genotypes and to avoid conducting experiments during the 2021 COVID-19 lockdowns in Melbourne.

### Lab experiment manipulating temperature and diet

In January 2022, we conducted a manipulative laboratory experiment with the five mass-bred populations to test how combined thermal and nutritional treatments influence clinal patterns in tolerance traits and body size. We picked eggs from all populations into six treatments, which included three constant temperatures (13, 25 and 29°C) and two diets (standard diet and a low-calorie 25% dilution diet). The 25°C condition served as the laboratory-acclimated control, while 13°C and 29°C represented the cold and hot extremes that larvae encounter in natural populations during winter and summer, respectively. The low-calorie diet was chosen to simulate reduced food availability predicted under climate change (Chakraborty et al. 2023). For each population, approximately 200 adult flies were placed in two laying pots with a thin layer of food at the bottom and allowed to lay eggs for six hours. We then transferred 20 eggs into each vial (n=36-42 vials/population/treatment; **Table S2**) containing 7mL of either standard or low-calorie (25% dilution) food and allowed larvae to develop at their respective treatment temperatures. To manage replication, the experiment was divided into two blocks, with adults emerging between January 21-28 (block 1) and March 1-8 (block 2).

For each block, we conducted 2-3 rounds of egg-picking for each treatment over 4-5 days to maximize overlap in eclosion across treatments (n=14-21 vials/population/treatment/block). We altered egg-picking times based on pilot experiments to ensure that adults would emerge simultaneously so that they could be randomised into the same thermal tolerance assays (**Table S2**). We then placed eggs into 36-42 vials per population per treatment combination, split across 5-6 rounds of egg-picking over the two experimental blocks (**Table S2**). In total, 2912 vials were used (n=455-546 vials per treatment).

When adults started eclosing, we collected them daily in separate vials for each treatment to ensure uniform age across treatments. We then kept flies at their experimental conditions until they matured and mated. In pilot experiments, we found that flies emerging at 13°C would not mature if kept at 13°C, so those collected from 13°C were kept at 16°C to mature in time for the experiment (seven days until egg-laying).

### Tolerance assays

We measured females as their body size and thermal tolerance are more sensitive to nutritional and temperature variation (Kutz et al. 2019; Chakraborty et al. 2020), and because sex-specific clinal trends were not the focus of our study. We collected females using light CO_2_ anesthesia, allowed them to recover for 48 hours, and then assayed them for cold, heat and desiccation tolerance following established methods (Hoffmann et al. 2002; Lasne et al. 2018). For each tolerance trait, we measured 30 females per population per treatment combination. For heat tolerance, we measured heat knockdown by placing individual females in sealed 5mL glass vials that we submerged in a 39°C water bath and then recorded the time it took each female to cease moving. For cold tolerance, we measured chill-coma recovery by placing individual females in sealed 1.5mL plastic tubes submerged in a 0°C water bath for four hours and then recorded the time they took to stand up at room temperature. For desiccation tolerance, we placed individual females in 5mL glass vials covered with mesh for airflow, which we placed within sealed tanks containing silica gel that maintained <5% relative humidity, and recorded the time to cessation of movement. For the tolerance traits, we were unable to measure all individual females in a single assay, and so we conducted 3-4 assays per block for each trait. Each assay included replicates of all populations in all treatment temperatures and diets. To measure body size, we quantified wing size by mounting the right wing of 30 females in SH solution (70% ethanol, 30% glycerol) on microscope slides, photographing them using a Leica M60 stereo microscope (Leica, Heerbrugg, Switzerland), and using ImageJ to measure 10 vein intersection landmarks (**Fig. S1**), which we used to estimate wing centroid size.

### Statistical analyses

We used *R* (v.4.4.1) statistical software for all analyses (R Core Team 2024). Using the package *glmmTMB* (Brooks et al. 2017), we implemented linear mixed models to test for clinal patterns in the four traits (thermal and desiccation tolerance, and wing size), and whether these patterns changed depending on treatment temperature and diet. We checked model assumptions using *DHARMa* (Hartig 2022). We tested each trait independently and included latitude (continuous trait) and treatment temperature and diet as the fixed effects, as well as their three-way interaction. We also included block as a fixed effect and assay (within block) as the only random effect. We then used *emmeans* (Lenth 2016) to obtain the marginal means for each population under each temperature and diet combination. While a significant latitude term would provide evidence for a latitudinal trend in a given trait, significant treatment×latitude terms would suggest that the clinal trends change depending on the treatment.

To test for clinal patterns in plasticity, we used the marginal means to quantify plasticity as the change in trait value from 25°C. We calculated plasticity separately for the two diets, meaning that we focus on plasticity in response to temperature. This was because we found stronger effects of temperature than diet, but the results did not change when we used only the 25°C, standard diet to calculate plasticity. We then used a linear model to test for significant clinal trends in plasticity, and significant effects of treatment temperature on clinal patterns.

## Results

### Clinal patterns in tolerance traits and body size change more with temperature than diet

We found no significant diet×temperature×latitude interactions for any trait (**Table 1**), suggesting that latitudinal trends did not change across all combinations of diet and temperature. Diet×latitude was only significant for cold tolerance, whereas temperature×latitude interactions were significant for all traits (**Table 1**). The strength of clinal patterns therefore depended more on temperature than diet. In general, we found evidence of latitudinal clines at 25°C whereby populations from higher latitudes showed higher cold tolerance, lower heat tolerance and larger wing (body) size, but no latitudinal cline for desiccation resistance (**Fig. 2**).

**Table 1.** ANOVA summaries for testing the stability of latitudinal clines for the four traits across the diet and temperature treatments. Terms that are significant at P<0.05 are denoted in bold.

| Term | Df | (a) Cold tolerance |  | (b) Heat tolerance |  | (c) Desiccation tolerance |  | (d) Wing centroid size |  |
| --- | --- | --- | --- | --- | --- | --- | --- | --- | --- |
| | | $\chi^2$ | P | $\chi^2$ | P | $\chi^2$ | P | $\chi^2$ | P |
| Intercept | 1 | 875.4 | **<0.001** | 1030.1 | **<0.001** | 1510.6 | **<0.001** | 63836.1 | **<0.001** |
| Latitude | 1 | 31.8 | **<0.001** | 52.5 | **<0.001** | 0.1 | 0.721 | 568.9 | **<0.001** |
| Temperature | 2 | 161.0 | **<0.001** | 261.0 | **<0.001** | 9.0 | **0.011** | 221.6 | **<0.001** |
| Diet | 1 | 7.5 | **0.006** | 0.1 | 0.748 | 34.0 | **<0.001** | 2.5 | 0.113 |
| Block | 1 | 33.9 | **<0.001** | 1.1 | 0.301 | 4.2 | **0.041** | - |  |
| Latitude × Temperature | 2 | 9.5 | **0.009** | 30.2 | **<0.001** | 10.8 | **0.005** | 10.7 | **0.005** |
| Latitude × Diet | 1 | 5.3 | **0.022** | 1.2 | 0.283 | 2.2 | 0.140 | 0.1 | 0.712 |
| Temperature × Diet | 2 | 1.8 | 0.405 | 0.5 | 0.794 | 0.7 | 0.701 | 2.7 | 0.264 |
| Latitude × Temperature × Diet | 2 | 1.6 | 0.450 | 0.1 | 0.970 | 0.6 | 0.748 | 4.7 | 0.097 |

**Fig. 2.**
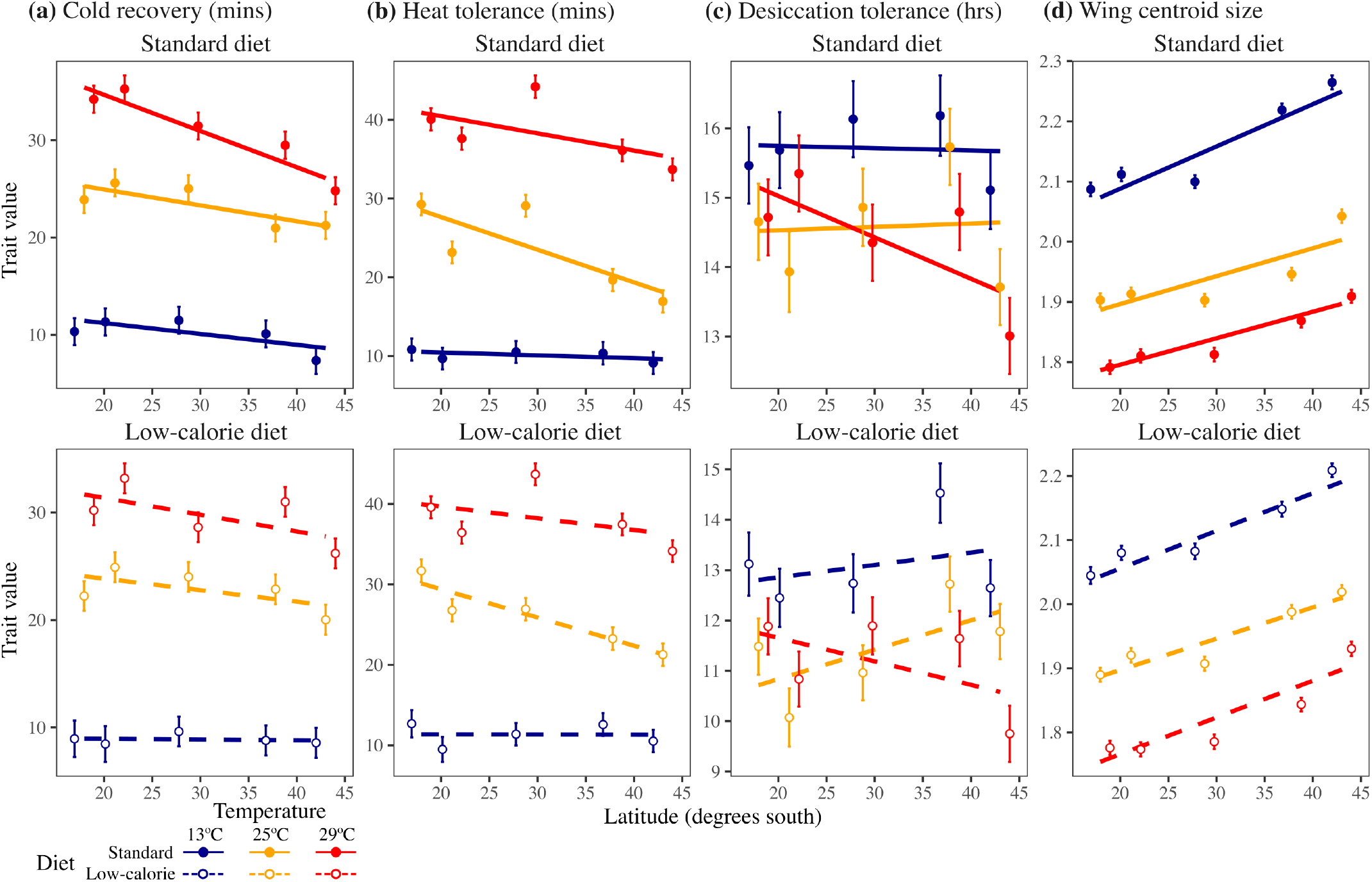
Testing for latitudinal clines in **(a)** cold tolerance, **(b)** heat tolerance, **(c)** desiccation tolerance, and wing size. Circles and credible intervals represent the mean (±1SE). Colours depict the different temperature treatments. Closed circles and solid lines represent the standard diet (upper panels), open circles and dashed lines the low-calorie diet (lower panels).

For cold tolerance, latitudinal clines were twice as strong at the standard diet (marginal slope=-0.22±0.04 mins per degree latitude [hereafter, mins/latitude]) compared to the low-calorie diet (marginal slope=- 0.09±0.04 mins/latitude), with the strongest clinal pattern present at the warmest treatment (**Fig. 2a; Table S3**). For heat tolerance, we found the strongest latitudinal cline at 25°C (marginal slope =- 0.38±0.05 mins/latitude), with a weaker clinal pattern at the warmest treatment (marginal slope for 29°C=-0.18±0.05 mins/latitude) and no latitudinal cline at the cold temperature (marginal slope for 13°C=-0.02±0.05 mins/latitude; **Fig. 2b; Table S3**). Desiccation tolerance showed weak positive trends across latitude for the cold and 25°C temperatures (marginal slope for 13°C=0.01±0.02 hrs/latitude, 25°C=0.03±0.02 hrs/latitude), suggesting that higher latitudes showed slightly greater desiccation tolerance. By contrast, desiccation tolerance was greater at lower latitudes in the warmest treatment (marginal slope for 29°C=-0.05±0.02 hrs/latitude), suggesting that lower latitudes showed greater desiccation tolerance at the warmest treatment (**Fig. 2c**). Body size increased at higher latitudes for all treatments (**Fig. 2d**), with the coldest treatment showing the strongest latitudinal cline (marginal slope for 13°C=0.006±0.0004 centroid size/latitude) compared to the warmer temperatures (marginal slope for 25°C=0.005±0.0004, 29°C=0.005±0.0004 size/latitude).

### Clinal patterns in plasticity are trait-specific and depend on temperature

Only plasticity in heat tolerance and desiccation tolerance showed significant latitudinal clines (**Fig. 3; Table 2**). For plasticity in heat tolerance, we found a significant latitude×temperature interaction (**Table 2b**), created by higher latitude populations showing greater increases in heat tolerance at 29°C (marginal slope=0.018±0.003) compared to 13°C (marginal slope=0.006±0.003; **Fig. 3b**). This meant that higher latitude populations showed greater plasticity, seen as a greater increase in heat tolerance at 29°C, and a smaller decrease in heat tolerance at 13°C (**Fig. 3b**). For plasticity in desiccation tolerance, populations from lower latitudes showed an increase in desiccation tolerance at both hot and cold temperatures relative to 25°C (**Fig. 3c**). By contrast, populations from higher latitudes showed lower plasticity as smaller increases in desiccation tolerance at 13°C (marginal slope=-0.002±0.002), but a large reduction in desiccation tolerance at 29°C (marginal slope=-0.007±0.002; **Fig. 3c**). Plasticity in body size and cold tolerance showed no significant latitudinal trends.

**Table 2.** ANOVA summaries for testing the stability of latitudinal clines in plasticity across temperatures (i.e., plastic change in trait from 25°C to each other temperature). Terms that are significant at P<0.05 are denoted in bold.

| Term | Df | (a) Cold tolerance |  |  | (b) Heat tolerance |  |  | (c) Desiccation tolerance |  |  | (d) Wing centroid size |  |  |
| --- | --- | --- | --- | --- | --- | --- | --- | --- | --- | --- | --- | --- | --- |
|  |  | SS | F | P | SS | F | P | SS | F | P | SS<br>(×10) | F | P |
| Intercept | 1 | 0.011 | 2.24 | 0.161 | 0.224 | 31.76 | **<0.001** | 0.064 | 22.88 | **<0.001** | 0.001 | 0.51 | 0.490 |
| Latitude | 1 | 0.007 | 1.37 | 0.264 | 0.263 | 37.27 | **<0.001** | 0.036 | 12.97 | **0.004** | 0.004 | 2.75 | 0.123 |
| Temperature | 1 | 0.48 | 96.44 | **<0.001** | 0.302 | 42.71 | **<0.001** | 0.000 | 0.02 | 0.883 | 0.101 | 69.89 | **<0.001** |
| Diet | 1 | 0.014 | 2.79 | 0.121 | 0.000 | 0.06 | 0.807 | 0.011 | 3.74 | 0.077 | 0.001 | 0.77 | 0.399 |
| Latitude × Temperature | 1 | 0.006 | 1.19 | 0.298 | 0.057 | 8.07 | **0.015** | 0.010 | 3.44 | 0.088 | 0.000 | 0.34 | 0.568 |
| Latitude × Diet | 1 | 0.010 | 2.11 | 0.172 | 0.007 | 1.06 | 0.324 | 0.007 | 2.49 | 0.141 | 0.000 | 0.01 | 0.908 |
| Temperature × Diet | 1 | 0.000 | 0.07 | 0.794 | 0.000 | 0.01 | 0.941 | 0.000 | 0.07 | 0.800 | 0.001 | 0.43 | 0.526 |
| Latitude × Temperature × Diet | 1 | 0.001 | 0.15 | 0.701 | 0.005 | 0.71 | 0.415 | 0.000 | 0.06 | 0.812 | 0.002 | 1.21 | 0.293 |
| Residual | 12 | 0.060 |  |  | 0.085 |  |  | 0.034 |  |  | 0.017 |  |  |

**Fig. 3.**
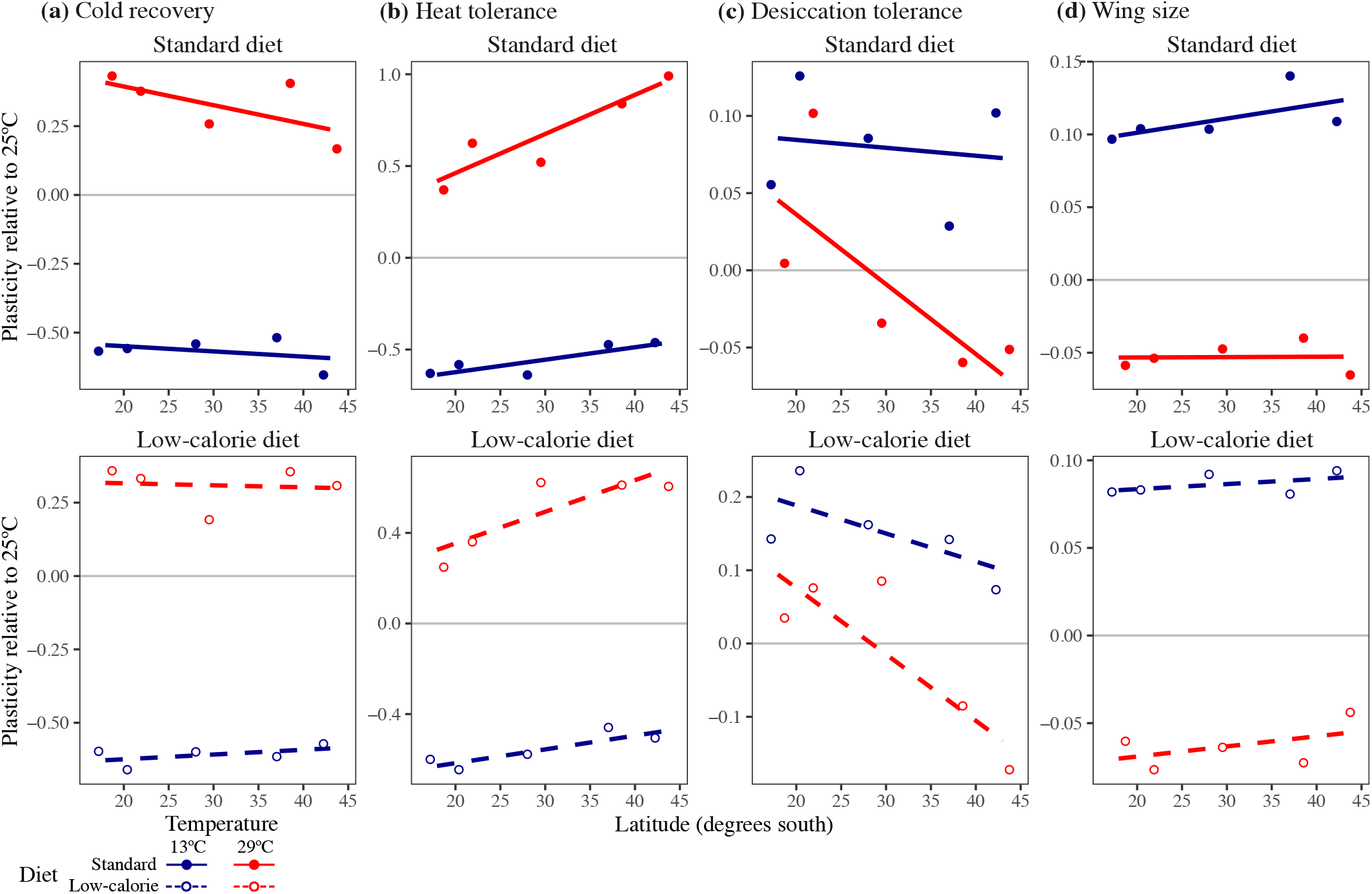
Testing for latitudinal clines in plasticity for **(a)** cold recovery, **(b)** heat tolerance, **(c)** desiccation tolerance, and **(d)** wing size. Colours depict the different temperature treatments. Closed circles and solid lines represent the standard diet (upper panels), open circles and dashed lines the low-calorie diet (lower panels). Plasticity values above and below zero increase and decrease the trait from 25°C, respectively.

## Discussion

Studies of latitudinal clines have demonstrated adaptation across broad climate gradients, but rarely consider whether patterns of adaptation remain stable across environments. Here, we tested how the strength of latitudinal clines in tolerance traits and their plasticity changed across combinations of diet and temperature treatments. We found that temperature influenced the strength of clinal patterns for all traits, while diet only influenced clinal variation in cold tolerance (**Table 1**). The hot treatment was associated with stronger latitudinal trends for cold and desiccation tolerance, but weaker trends for heat tolerance, whereas stronger clines in body size emerged at the coldest temperature (**Fig. 2**). Latitudinal clines generally followed previously reported patterns: populations from lower latitudes showed lower cold tolerance, higher heat tolerance and smaller wing sizes (James et al. 1995; Hoffmann et al. 2002), but surprisingly, higher desiccation tolerance (but only under hot conditions). Temperature also determined the strength of latitudinal clines for plasticity in heat tolerance and desiccation tolerance (**Fig. 3b-c**). In the hot treatment, higher latitude populations showed greater plasticity as a greater increase in heat tolerance and as a stronger reduction in desiccation tolerance. These results provide evidence that the strength of latitudinal clines is sensitive to variation in temperature, and that environmental change is likely to affect populations adapted across latitude differently.

### Treatment temperature influenced the strength of latitudinal clines more than diet

Treatment temperature created variation in the strength of latitudinal clines for all traits, whereas the low-calorie diet had little effect on the strength of latitudinal clines. Only cold recovery showed significant latitude×temperature and latitude×diet interactions, which supports evidence that cold recovery exhibits strong genotype-by-environment interactions (Ørsted et al. 2019; Littler et al. 2021). Our results therefore suggest that changes in the availability of food are less likely than temperature to override signals of climatic adaptation. This is consistent with a study of *Sepsis* flies that showed temperature but not diet influenced latitudinal divergence in genetic variance underlying development rate (Berger et al. 2013). However, a study that manipulated the protein and carbohydrate availability in diet treatments for three populations of *D. melanogaster* along a latitudinal cline found that populations varied in plasticity in response to diet, but in a way that was idiosyncratic for populations across latitude (Chakraborty et al. 2020). This is consistent with another study that found populations of *D. melanogaster* along latitude varied idiosyncratically in their immune system activation, which erased the latitudinal cline in heat tolerance (Hector et al. 2020). Given that food composition and availability is unlikely to show a similar latitudinal gradient to temperature, it is possible that adaptation to nutrition along the east coast of Australia will be region specific, creating patterns of local adaptation without a latitudinal cline.

The strength of latitudinal clines changed across treatment temperatures in a manner that was trait-dependent. For cold recovery, we observed the strongest latitudinal cline at the warmest temperature (**Fig. 2a**), which was because lower latitude populations took longer to recover from a chill coma after developing in the hot treatment. Low latitude populations therefore showed lower cold tolerance at the hot treatment compared to higher latitude populations. This is consistent with evidence that populations from low latitudes are closer to their higher thermal limits and are likely to struggle with other stressors, such as rapid changes in temperature (Kellermann et al. 2012; van Heerwaarden et al. 2016). Body (wing) size showed the strongest latitudinal cline in the cold treatment (**Fig. 2d**), suggesting that higher latitude populations increased their body size in the cold temperature marginally more than the lower latitude populations (**Fig. 3d**). Both heat tolerance and desiccation tolerance showed strong clinal patterns that changed with temperature (**Fig. 2b-c**). Below we discuss the effects of plasticity on determining the strength of latitudinal clines in heat tolerance and desiccation tolerance.

### The stability of latitudinal clines was linked to plasticity, but patterns were trait-dependent

Our observed latitudinal clines in plasticity were likely responsible for changes in the strength of clinal patterns across temperature, particularly for heat and desiccation tolerance. While strong patterns of adaptation (i.e., strong latitudinal clines) were found at 25°C for heat tolerance (**Fig. 2b**), the weaker clinal trends found at the warmest treatment highlights that assessing clinal trends only under benign laboratory-acclimated conditions could underestimate the potential for higher latitude populations to maintain resilience in response to ongoing climate change. This is because higher latitude populations showed the greatest plastic increase in heat tolerance at 29°C (**Fig. 3b**). Therefore, supporting both hypotheses (climate variability and plasticity trade-off), populations from higher latitudes showed lower heat tolerance, but greater plasticity in heat tolerance. These results also suggest that trade-offs could be more likely to emerge at warmer temperatures, which will be important to consider for predicting population responses to climate change (van Heerwaarden and Kellermann 2020; Bogan et al. 2024; van Heerwaarden et al. 2024).

Neither cold or desiccation tolerance supported the climate variability or trade-off hypotheses. Populations from higher latitudes showed faster cold recovery and lower plasticity (smaller increase) in the hot treatment (**Fig. 3a**). More variable lower temperatures at higher latitudes could therefore increase cold tolerance without compromising adaptive plasticity (cold recovery was maintained at high temperature). Desiccation tolerance showed zero or weak positive latitudinal clines in cold or 25°C temperatures, but a strong negative latitudinal cline in the hottest treatment (**Fig. 2c**). Higher latitude populations were therefore the least desiccation tolerant at 29°C, which was associated the greatest plastic decrease in desiccation tolerance (**Fig. 3c**). Higher latitude populations therefore could not maintain desiccation tolerance when exposed to warmer temperatures, despite showing the strongest ability to increase heat tolerance. By contrast, the low latitude populations showed lower plasticity in heat tolerance but a better ability to maintain desiccation resistance at 29°C. Adapting to more constant high temperatures at low latitudes could therefore preadapt populations for desiccation tolerance at high temperatures. Together, these results suggest that locally adapted populations show different trade-offs among traits that could determine responses to environmental change (Hoffmann 1990; Hoffmann 1991; Terblanche and Kleynhans 2009; Kellermann et al. 2020; van Heerwaarden and Kellermann 2020). Determining how interactions among traits and their plasticity determines adaptive responses to climate change therefore remains an ongoing challenge for predicting the resilience of natural populations.

### Predicting population responses to climate change

Our results support the theory that considering how locally adapted populations vary in plasticity will be key to accurately predicting their response to global change (Chevin et al. 2013; Valladares et al. 2014). Populations from higher latitudes showed greater plasticity in heat tolerance, which is consistent with studies on *Ascaphus* frogs demonstrating populations from more variable environments showed greater plasticity (Cicchino et al. 2024), and on *Drosophila* where a temperate population of *Drosophila* showed greater plasticity than a tropical population (van Heerwaarden and Sgrò 2017). Our results therefore support growing evidence that accurately predicting the capacity for climate adaptation requires considering how populations (within a species) evolve plasticity as they locally adapt to environmental heterogeneity (Overgaard et al. 2011). By contrast, meta-analyses comparing variation among species found little support for the climate variability hypothesis, suggesting that plasticity has a limited capacity for helping ectotherms to cope with climate change (Gunderson and Stillman 2015; MacLean et al. 2019). However, we show that considering variation among species without accounting for variation within species could overlook adaptive variation in plasticity that could be crucial for climate adaptation.

### Caveats

Given that changes in climate across latitude involve coordinated changes in humidity, temperature and their variability (**Table S1**), it is difficult to distinguish adaptation to temperature from the other autocorrelated environmental variables. Assaying populations from a variety of thermal environments can provide a contrast to show how both variability and predictability in temperature drives adaptation in thermal tolerance (Walter et al. 2025a). Also considering how much variation among genotypes and subpopulations within each latitudinal location will be important for predicting the potential to adapt to rapid environmental change (Hoffmann et al. 2001), rather than relying on latitudinal clines. Finally, it is surprising that diet had little impact on clinal patterns, and could suggest that our low-calorie diet, which represents one scenario of environmental change, does not affect patterns of adaptation. Because diet is unlikely to change gradually across latitude like temperature, it is possible that local adaptation to diet depends on the availability of different types of food. While diet effects can be idiosyncratic and difficult to predict for locally adapted populations (Chakraborty et al. 2020), changes in the type of nutrition (rather than just quantity, as in our study) affects life history traits, body size and metabolic responses to warmer temperatures (Kutz et al. 2019; Alton et al. 2020), and so further work is needed to disentangle how variation in diet could influence patterns of adaptation.

### Conclusions

Our results reveal that predicting population responses to climate change based only on a single environmental condition can be inaccurate. The strength of latitudinal clines in tolerance traits and body size changed dramatically with temperature, but was not strongly influenced by diet treatment. Populations at higher latitudes showed lower heat tolerance but higher plasticity in heat tolerance, supporting the climate variability hypothesis as well as a trade-off between basal heat tolerance and plasticity. However, plasticity in cold and desiccation tolerance was not consistently greater for high latitude populations, suggesting that adaptive plasticity is trait-dependent and could be determined by trade-offs among traits. Together, our results show that accurately predicting population responses to climate change requires understanding how populations locally adapt to variation in environmental heterogeneity, how this determines the evolution of plasticity, and the consequences for coping with climate change.

