## Supplementary material for "Variation in temperature but not diet determines the stability of latitudinal clines in tolerance traits and their plasticity"

**Contents**

**Table S1** Sampling locations

**Table S2** Developmental time of each treatment

**Fig. S1** Wing landmarks

**Table S3** Slope estimates for the latitudinal clines

**Table S1:** Population locations sampled from the cline and across Victoria.

| Code | State | Location | Sampling location | Latitude | Longitude | Number of lines | Mean annual rainfall (mm) | Mean diurnal range (°C) | Temperature seasonality (sd ×100) |
|---|---|---|---|---|---|---|---|---|---|
| INI | Qld | Innisfail | Sellars Bananas | -17.9414 | 146.0553 | 40 | 2844 | 7.9 | 282 |
| MAC | Qld | Mackay | FoodPac Fruit | -21.1363 | 148.5098 | 47 | 1464 | 9.6 | 374 |
| BAL | NSW | Ballina | Southern Cross Botanicals | -28.7670 | 153.5361 | 19 | 1782 | 10 | 362 |
| TAS | Tas | Huonville | Willie Smith's Organic Cider | -42.9960 | 147.0718 | 29 | 1167 | 10.1 | 394 |
| RY | Vic | Yarra Valley | Raynor's Orchard | -37.7924 | 145.5512 | 44 | 751 | 10.5 | 327 |

**Table S2:** Developmental time of each treatment

| Temperature | Diet | Time to eclosion | Block 1 | Block 2 |
|---|---|---|---|---|
| 13°C | Control | 36-38 days | 3 egg-picks<br>21 vials/population | 3 egg-picks<br>21 vials/population |
| | 25% dilution | 42-46 days | 3 egg-picks<br>21 vials/population | 3 egg-picks<br>21 vials/population |
| 25°C | Control | 9-10 days | 2 egg-picks<br>14 vials/population | 3 egg-picks<br>21 vials/population |
| | 25% dilution | 10-11 days | 2 egg-picks<br>14 vials/population | 3 egg-picks<br>21 vials/population |
| 29°C | Control | 7-8 days | 2 egg-picks<br>14 vials/population | 3 egg-picks<br>21 vials/population |
| | 25% dilution | 8-9 days | 2 egg-picks<br>14 vials/population | 3 egg-picks<br>21 vials/population |

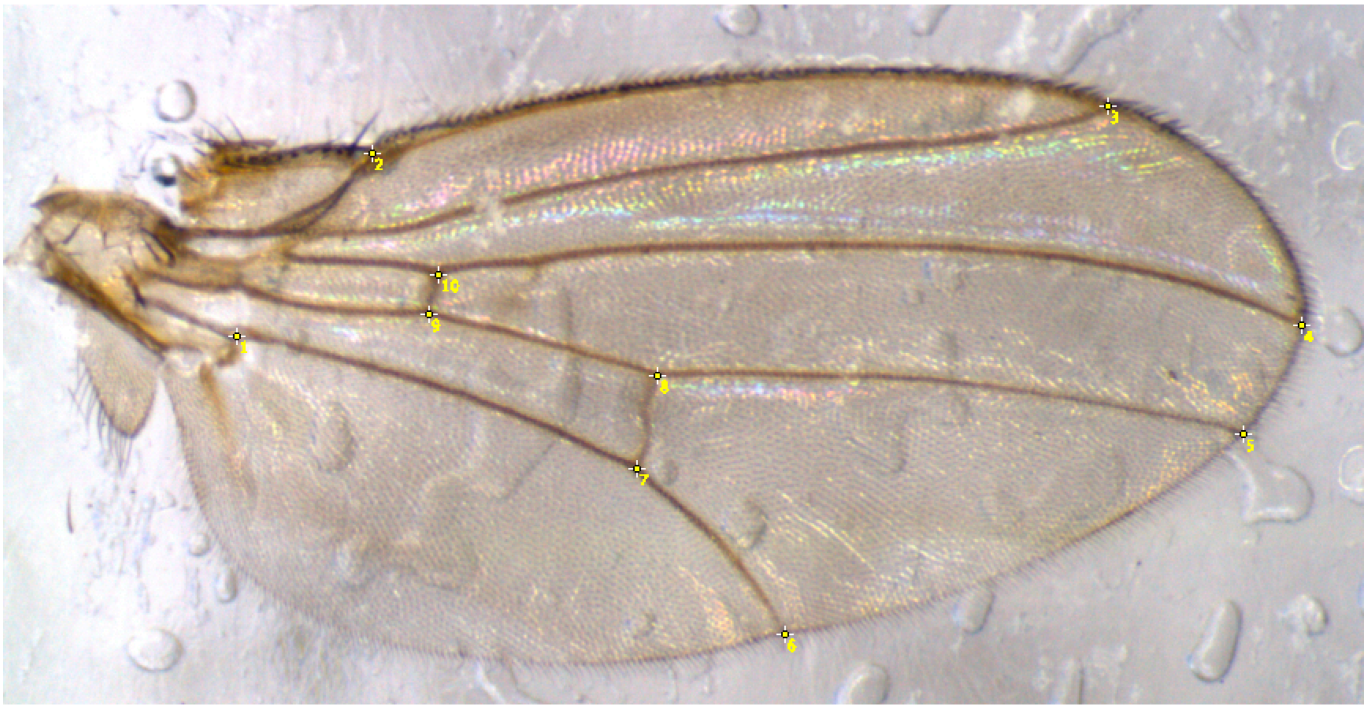

**Fig. S1** Wing landmarks used to estimate centroid size.

**Table S3** Slope estimates for latitudinal clines in **(a)** tolerance traits, and **(b)** their plasticity.

**(a)** Tolerance traits

| Temperature | Diet | Latitude slope | SE |
|---|---|---|---|
| 13°C | Standard | -0.112 | 0.0655 |
| 25°C | | -0.164 | 0.0655 |
| 29°C | | -0.373 | 0.0657 |
| 13°C | 25% | -0.01 | 0.0724 |
| 25°C | | -0.107 | 0.0655 |
| 29°C | | -0.156 | 0.0655 |
| 13°C | Standard | -0.037 | 0.0651 |
| 25°C | | -0.415 | 0.0649 |
| 29°C | | -0.219 | 0.0653 |
| 13°C | 25% | 0.002 | 0.0708 |
| 25°C | | -0.35 | 0.0653 |
| 29°C | | -0.149 | 0.0627 |
| 13°C | Standard | -0.004 | 0.0265 |
| 25°C | | 0.004 | 0.0263 |
| 29°C | | -0.06 | 0.0261 |
| 13°C | 25% | 0.024 | 0.0279 |
| 25°C | | 0.058 | 0.0265 |
| 29°C | | -0.045 | 0.0265 |
| 13°C | Standard | 0.007 | 0.0006 |
| 25°C | | 0.005 | 0.0006 |
| 29°C | | 0.004 | 0.0006 |
| 13°C | 25% | 0.006 | 0.0006 |
| 25°C | | 0.005 | 0.0005 |
| 29°C | | 0.006 | 0.0006 |

**(b)** Plasticity in tolerance traits

| Temperature | Diet | Latitude slope | SE |
|---|---|---|---|
| 13°C | Standard | -0.002 | 0.0033 |
| 29°C | | -0.007 | 0.0033 |
| 13°C | 25% | 0.002 | 0.0033 |
| 29°C | | -0.001 | 0.0033 |
| 13°C | Standard | 0.007 | 0.0039 |
| 29°C | | 0.021 | 0.0039 |
| 13°C | 25% | 0.006 | 0.0039 |
| 29°C | | 0.014 | 0.0039 |
| 13°C | Standard | -0.001 | 0.0025 |
| 29°C | | -0.005 | 0.0025 |
| 13°C | 25% | -0.004 | 0.0025 |
| 29°C | | -0.009 | 0.0025 |
| 13°C | Standard | 0.001 | 0.0006 |
| 29°C | | 0 | 0.0006 |
| 13°C | 25% | 0 | 0.0006 |
| 29°C | | 0.001 | 0.0006 |
